# NuRD chromatin remodeling is required to repair exogenous DSBs in the *Caenorhabditis elegans* germline

**DOI:** 10.1101/2024.09.14.613027

**Authors:** Deepshikha Ananthaswamy, Kelin Funes, Thiago Borges, Scott Roques, Nina Fassnacht, Sereen El Jamal, Paula M. Checchi, Teresa Wei-sy Lee

**Author notes:** contributed equally (Kelin Funes, Thiago Borges).

## Abstract

Organisms rely on coordinated networks of DNA repair pathways to protect genomes against toxic double-strand breaks (DSBs), particularly in germ cells. All repair mechanisms must successfully negotiate the local chromatin environment in order to access DNA. For example, nucleosomes can be repositioned by the highly conserved Nucleosome Remodeling and Deacetylase (NuRD) complex. In *Caenorhabditis elegans*, NuRD functions in the germline to repair DSBs – the loss of NuRD’s ATPase subunit, LET-418/CHD4, prevents DSB resolution and therefore reduces fertility. In this study, we challenge germlines with exogenous DNA damage to better understand NuRD’s role in repairing DSBs. We find that *let-418* mutants are sensitive to cisplatin and hydroxyurea: exposure to either mutagen impedes DSB repair, generates aneuploid oocytes, and reduces fertility and embryonic survival. These defects resemble those seen when the Fanconi anemia (FA) DNA repair pathway is compromised, and we find that LET-418’s activity is epistatic to that of the FA component FCD-2/FANCD2. We propose a model in which NuRD is recruited to the site of DNA lesions to remodel chromatin and allow access for FA pathway components. Together, these results implicate NuRD in the repair of both endogenous DSBs and exogenous DNA lesions to preserve genome integrity in developing germ cells.

**ARTICLE SUMMARY:** Preserving genome integrity in germ cells is critical for the survival of individuals and species. Our previous work shows that nucleosome remodeling plays an important role in repairing meiotic DNA damage. Here, we further challenge genomes with toxic DNA damaging agents to test the requirement for remodeling in the germline. We find that DNA damage accumulates in the absence of remodeling, which drastically reduces oocyte quality, and also show that the requirement for remodeling is epistatic to the Fanconi anemia DNA repair pathway. These findings demonstrate that local chromatin environments must be remodeled in response to DNA damage to maintain oocyte quality.

## INTRODUCTION

DNA damage poses a danger to all cells, but if left unrepaired in germ cells, it is deleterious for organismal survival. During meiosis, any DNA lesion that persists until chromosome segregation will generate an aneuploid gamete, which in turn causes infertility or embryonic lethality (Ceccaldi et al., 2016; Kim et al., 2016). The evolutionary pressure to protect genomes for future generations has generated multiple distinct, but overlapping, DNA repair pathways. These pathways handle both endogenous programmed double-strand breaks (DSBs) required for crossover recombination, along with any exogenous DNA damage incurred in the germline (Chapman et al., 2012; Gartner and Engebrecht, 2022; Keeney, 2007). Studies in organisms ranging from budding yeast to mammals have identified strong conservation among these repair pathways, which include error-free mechanisms like homologous recombination and error-prone mechanisms like non-homologous end joining (De Massy, 2013; Stinson and Loparo, 2021, 2021; Wilson et al., 1997). For example, the Fanconi anemia (FA) pathway is activated when replication forks collide with DNA interstrand crosslinks, which are one of the most toxic DNA lesions (Gartner and Engebrecht, 2022). Components of the FA pathway coordinate the excision of the crosslink and conversion to DSBs, which can then be resolved by the meiotic homologous recombination pathway (Adamo et al., 2010; Lachaud et al., 2016; Raghunandan et al., 2015; Schlacher et al., 2012). Defects in the FA pathway are linked to an elevated risk of cancer and cause chromosomal instability or meiotic catastrophe, hallmarks of the rare congenital disorder of Fanconi anemia (Auerbach, 2009, 1993; Ceccaldi et al., 2016; De Winter and Joenje, 2009; Nalepa and Clapp, 2018; Tischkowitz et al., 2003, 2004).

In eukaryotes, DNA repair must always happen in the context of chromatin, where DNA is wrapped around histone octamers to form nucleosomes (Allis and Jenuwein, 2016). Nucleosomes act as obstacles that control access to DNA, and therefore their spacing is regulated by nucleosome remodeling complexes (Becker and Workman, 2013). The highly conserved remodeling complex, <u>Nu</u>cleosome remodeling and <u>d</u>eacetylase (NuRD), includes ATPase remodelers from the <u>c</u>hromodomain <u>h</u>elicase <u>D</u>NA binding (CHD) family (Basta and Rauchman, 2015; Kehle, 1998; Tong et al., 1998; Wade et al., 1998; Woodage et al., 1997; Xue et al., 1998). In addition to CHD remodeling subunits, NuRD also consists of the histone deacetylases HDAC1/2, histone demethylase LSD1/KDM1A, and other components that allow NuRD to bind modified histones, methyl CpG, and other proteins (Basta and Rauchman, 2015; Wang et al., 2009; Whyte et al., 2012). Initially, NuRD’s primary role was thought to be transcriptional repression; but studies have since identified a more nuanced involvement in gene expression, along with other biological functions like DNA repair (Basta and Rauchman, 2015). The nematode *Caenorhabditis elegans* has two catalytic CHD paralogs associated with the core NuRD complex: CHD-3 (also known as Mi-2α or CHD3 in mammals) and LET-418 (Mi-2β or CHD4) (Passannante et al., 2010; von Zelewsky et al., 2000). Mutations in either the *chd-3* or the *let-418* genes severely reduce fertility due to high levels of persistent meiotic DSBs (McMurchy et al., 2017; Turcotte et al., 2018).

These studies highlight the importance of NuRD’s role in the germline for proper DNA repair – the loss of NuRD activity appears to shunt DSBs from error-free repair towards error-prone pathways like NHEJ. The genomic instability of *chd-3* or *let-418* mutants resembles those seen in mutants defective for the FA pathway (Rageul and Kim, 2020; Tsui and Crismani, 2019; Turcotte et al., 2018). One hallmark of mutations in FA components is an increased sensitivity to mutagens like cisplatin, which induces DNA interstrand crosslinks, or hydroxyurea, which stalls replication forks (Bailly and Gartner, 2011; Datta and Brosh Jr., 2019; Youds et al., 2009). Both of these lesions can generate DSBs if left unrepaired, which activate the S-phase checkpoint and subsequently delays cell cycle progression (Kessler and Yanowitz, 2014; Kim and Colaiácovo, 2015). Resolution of interstrand crosslinks by the FA pathway involves recognition and binding by the core FA complex, which then recruits FANCD2 to chromatin by mono-ubiquitination (Nakanishi et al., 2005; Niraj et al., 2019). Once at the DNA lesion, FANCD2 orchestrates the repair of the lesion by converting it to DSBs, which are then resolved by homologous recombination (Ceccaldi et al., 2016; Niraj et al., 2019).

NuRD is required for repairing DNA lesions in germ cells (McMurchy et al., 2017; Turcotte et al., 2018), but it is not clear how or whether nucleosome remodeling interacts with repair processes like the FA pathway. Here, we challenge the germline by inducing exogenous DNA damage to investigate the relationship between the FA pathway and NuRD’s catalytic component, LET-418/CHD4. *C. elegans* has homologs for many core FA components, including FCD-2/FANCD2 (Kim et al., 2018; Lee et al., 2010). One advantage of using *C. elegans* for this study is the germline’s spatiotemporal organization, which allows us to easily follow the effects of mutagen exposure at multiple downstream stages during oogenesis (Jaramillo-Lambert et al., 2007; Kessler and Yanowitz, 2014). We show that LET-418 activity is epistatic to FCD-2 and is necessary to repair germline DSBs and preserve fertility. These results support a model where NuRD generates a permissive local chromatin environment at the site of DNA damage to allow access to DSB repair machinery and therefore preserve gamete quality in the face of genotoxic stress.

## METHODS

### Strains and husbandry

All strains were cultured using standard methods (Brenner, 1974) at 20°C on 6-cm NGM agar plates spotted with OP50 *E. coli* grown overnight at 37°C in Luria Broth (LB). Gene and sequence information was downloaded from WormBase (Davis et al., 2022). The *syb6818* allele is a deletion of LET-418’s ATPase domain and removes amino acids 590-1114. The *syb6708* allele is a deletion of LET-418’s PHD finger domain and removes amino acids 256-365. Both domains were identified using an NCBI structure prediction and based on homology with human CHD4 (Farnung et al., 2020; Sayers et al., 2022). TER6 is a homozygous sterile strain, and was maintained over an *nT1* balancer chromosome by picking GFP-positive heterozygotes. The following strains were obtained from the *Caenorhabditis* Genetics Center or were generated by SunyBiotech (*syb* alleles, which were backcrossed five times to N2):

- N2: wild-type (Bristol isolate)
- MT14390: *let-418 (n3536) V*
- NB105: *fcd-2 (tm1298) IV*
- TG1660: *xpf-1 (tm2842) II*
- TER14: *fcd-2 (tm1298) IV; let-418 (n3536) V*
- TER6: *let-418 (syb6818)/nT1 [qls51] IV:V*
- TER10: *let-418 (syb6708) V*

### Mutagen exposure

All exposure assays were performed on hermaphrodites cultured at 20°C on 3-cm NGM agar plates. Plates were first spotted with OP50 *E. coli*, then treated by spreading 250 μL of either 400 μM cisplatin (Sigma Aldrich) or 10 mM hydroxyurea (Thermo Scientific) onto agar at least 12 hours prior to experiment. Synchronized young adult hermaphrodites (12 hours post-L4 larval stage) were placed on mutagen plates for 24 hours, then moved to regular NGM plates to recover before assessing nuclei in the appropriate meiotic stage; the timing for each assay is described below (Jaramillo-Lambert et al., 2007; Kessler and Yanowitz, 2014).

### Immunofluorescence and image processing

To assess RAD-51 foci in the mitotic region after mitotic exposure, exposed and control hermaphrodites were allowed to recover on NGM plates for four hours before dissection to produce better gonad extrusion. To assess RAD-51 foci in pachytene after mitotic exposure, hermaphrodites were allowed to recover for 30-40 hours, allowing nuclei to progress to mid-to late-pachytene. Gonad dissection, fixation, and immunostaining were performed as previously described (Ananthaswamy et al., 2022). In brief, gonads were dissected using 25-gauge syringe needles, fixed in 0.8% paraformaldehyde (Electron Microscopy Services) for five minutes at room temperature, subjected to freeze crack, and dehydrated in 100% methanol for one minute at -20°C. Slides were washed three times with PBS + 0.1% Tween-20 (Thermo Scientific) prior to overnight incubation at room temperature with primary antibody. Slides were washed three times before incubation with secondary antibody for four hours at room temperature. Samples were mounted in VectaShield (Vector Laboratories) with 2 μg/ml DAPI (Thermo Life) before sealing with nail polish. Antibodies used were rabbit anti-RAD-51 at 1:5000 (a gift from Diana Libuda (Kurhanewicz et al., 2020)) and donkey anti-rabbit AlexaFluor594 at 1:400 (Invitrogen).

Immunofluorescence images were collected at 63x as Z-stacks with 0.25 μm intervals using a Zeiss Axio Imager M2 microscope with Zen 3.8 software (Zeiss). All representative images shown are max projections after processing in FIJI software (Schindelin et al., 2012). Unprocessed Z-stacks were used to quantify RAD-51 foci. Three biological replicates were performed, with three to five gonads imaged per condition in each replicate. Only one gonad arm was analyzed from each hermaphrodite. In the mitotic region, all nuclei were scored for RAD-51 foci. In pachytene, only nuclei in the top row were scored for RAD-51 foci; pachytene regions were determined by counting the number of rows after the transition zone and before diplotene, and dividing into equal thirds. Some nuclei had a distinctive apoptotic appearance with collapsed, DAPI-bright chromosomes and an excess of RAD-51 staining; these were included in the total nuclei number, but not scored for RAD-51 foci. All statistical analyses were performed in Prism 8 (GraphPad), with *P*-values determined using a Kruskal-Wallis non-parametric test after a Dunn’s post-hoc test used to account for pairwise comparisons.

### Analysis of diakinesis DAPI bodies

Following exposure to cisplatin or hydroxyurea, hermaphrodites were allowed to recover for 60 hours, allowing mitotic oocyte nuclei to progress through meiosis to diakinesis. Gonads were dissected as previously described (Ananthaswamy et al., 2022) with the following modifications: the primary and secondary antibody incubation steps were skipped, and slides were washed at least three times in PBS + 0.1% Tween-20 (Thermo Scientific) before mounting in SlowFade Diamond (Molecular Probes) with 2 μg/ml DAPI (Thermo Life). Images were collected at 63x as Z-stacks with 0.25 μm intervals captured on Zeiss Axio Imager M2 using Zen 3.8 software (Zeiss). All representative images shown are max projections after processing in FIJI (Schindelin et al., 2012). Unprocessed z-stacks were used to quantify DAPI-stained bodies in the -1 to the -4 oocyte from the spermatheca. Three biological replicates were performed, with each condition consisting of a minimum of 100 nuclei from at least 27 hermaphrodites. All statistical analyses were performed in Prism 10 (GraphPad), with p-values determined using a Fisher’s exact test on a 2x5 contingency table.

### Embryonic survival – 4-hr laying period

After exposure to cisplatin, hydroxyurea, or nitrogen mustard, hermaphrodites were allowed to recover on NGM plates for 60 hours, allowing mitotic oocyte nuclei to progress through the gonad and be laid as fertilized embryos. Exposed mothers were then moved onto freshly spotted 3-cm NGM plates (five to ten mothers per plate) and allowed to lay for four hours before being removed from plates. The number of embryos per plate was immediately recorded after removal of mothers, and were scored as fertilized, unfertilized, or dead. Hatched progeny were counted as adults three days later. Percent survival was calculated by dividing the number of hatched progeny by the number of embryos, multiplied by 100. Each condition consists of progeny from 125-150 mothers, collected over three biological replicates. All statistical analyses were performed in Prism 8 (GraphPad), with p-values determined from a one-way ANOVA after a Šídák correction used to account for multiple comparisons.

### Embryonic survival – entire brood

Individual hermaphrodites were isolated on separate plates as L4 larvae and transferred every 12 hours until the end of their fertile period (usually 5-6 days). Embryos and unfertilized eggs were counted every 12 hours immediately after transfer of the mother. Total brood was calculated for each mother as all embryos laid during the fertile period. Hatched progeny were then counted two to three days later (when they were L4s or young adults). Percent survival was calculated by dividing the number of hatched progeny by the number of embryos laid and multiplied by 100. Each biological replicate includes the broods of five mothers scored for each genotype (i.e. five technical replicates). The following biological replicates were performed: seven of N2; six of *let-418 (n3536)*; four of *let-418 (syb6818)*; and four of *let-418 (syb6708)*. Broods were censored from analysis if the mother died of unnatural causes before the end of their fertile period (usually from vulva rupture or crawling off the agar). All statistical analyses were performed in Prism 8 (GraphPad), with p-values determined from a one-way ANOVA after a Šídák correction used to account for multiple comparisons.

## RESULTS

### LET-418 activity is required to repair mitotic DNA lesions in mitosis and meiosis

Mutants lacking both NuRD paralogs (CHD3 and CHD4) have severely compromised fertility, which makes downstream analyses difficult (Turcotte et al., 2018). Therefore, we focused only on the impact of LET-418/CHD4, reasoning that it plays a larger role in repairing germline DSBs because its loss causes more meiotic defects than that of CHD-3 (Turcotte et al., 2018). A previous study noted that at pachytene entry, *let-418* mutants have more DSBs than wild-type (Turcotte et al., 2018), leading us to wonder whether these DSBs represent unrepaired DNA damage acquired during replication. Because DSBs are rare in mitotic nuclei, we used cisplatin and hydroxyurea to generate exogenous DNA damage and further stress the DNA repair process. We examined *let-418 (n3536)* missense hypomorphic mutants and *fcd-2 (tm1298)* deletion mutants, which lack the FA pathway component FCD-2/FANCD2. We capitalized on the fact that the *C. elegans* germline is organized in a spatiotemporal manner, with nuclei undergoing mitosis at the distal tip and moving proximally as they proceed through meiosis. The timing of how nuclei progress through mitotic replication, meiosis, oocyte maturation, and fertilization has been extensively characterized (Jaramillo-Lambert et al., 2007; Kessler and Yanowitz, 2014). We therefore estimated the timing after exposure to capture the impact of DNA damage incurred during mitosis.

Mitotic zone DSBs were assessed using an antibody for the RecA recombinase RAD-51, a common cytological marker for DSBs (Alpi et al., 2003; Colaiácovo et al., 2003; Kurhanewicz et al., 2020). Hermaphrodites were cultured on either cisplatin or hydroxyurea for twenty hours, allowed to recover for four hours, and dissected for immunofluorescence (Jaramillo-Lambert et al., 2007; Kessler and Yanowitz, 2014). Consistent with prior reports, unexposed wild-type animals accumulated few RAD-51 foci in the mitotic zone (Fig. 1A, C, Fig. 2A, C, and Table S1) (Alpi et al., 2003; Turcotte et al., 2018), which was also true for all mutants examined, with foci in less than 8% of nuclei. Unexposed *let-418* single mutants had a modest but significant increase in foci compared to wild-type animals (*P* < 0.0001, Kruskal-Wallis test), but neither *fcd-2* single mutants nor *fcd-2; let-418* double mutants had more foci than wild-type (Fig. 1A, *P* > 0.08 for both comparisons, Kruskal-Wallis test). Unexposed *fcd-2; let-418* double mutants were not more affected than either single mutant (*P* > 0.8 for both comparisons, Kruskal-Wallis test). We note that, although the average number of foci did not vary much between genotypes, we occasionally observed nuclei with two or three foci in *let-418* single mutants (0.8% of nuclei) and *fcd-2; let-418* double mutants (0.7%), which were rarely seen in wild-type germlines (0.2%) (Fig. 1A and Table S1).

**Figure 1:**
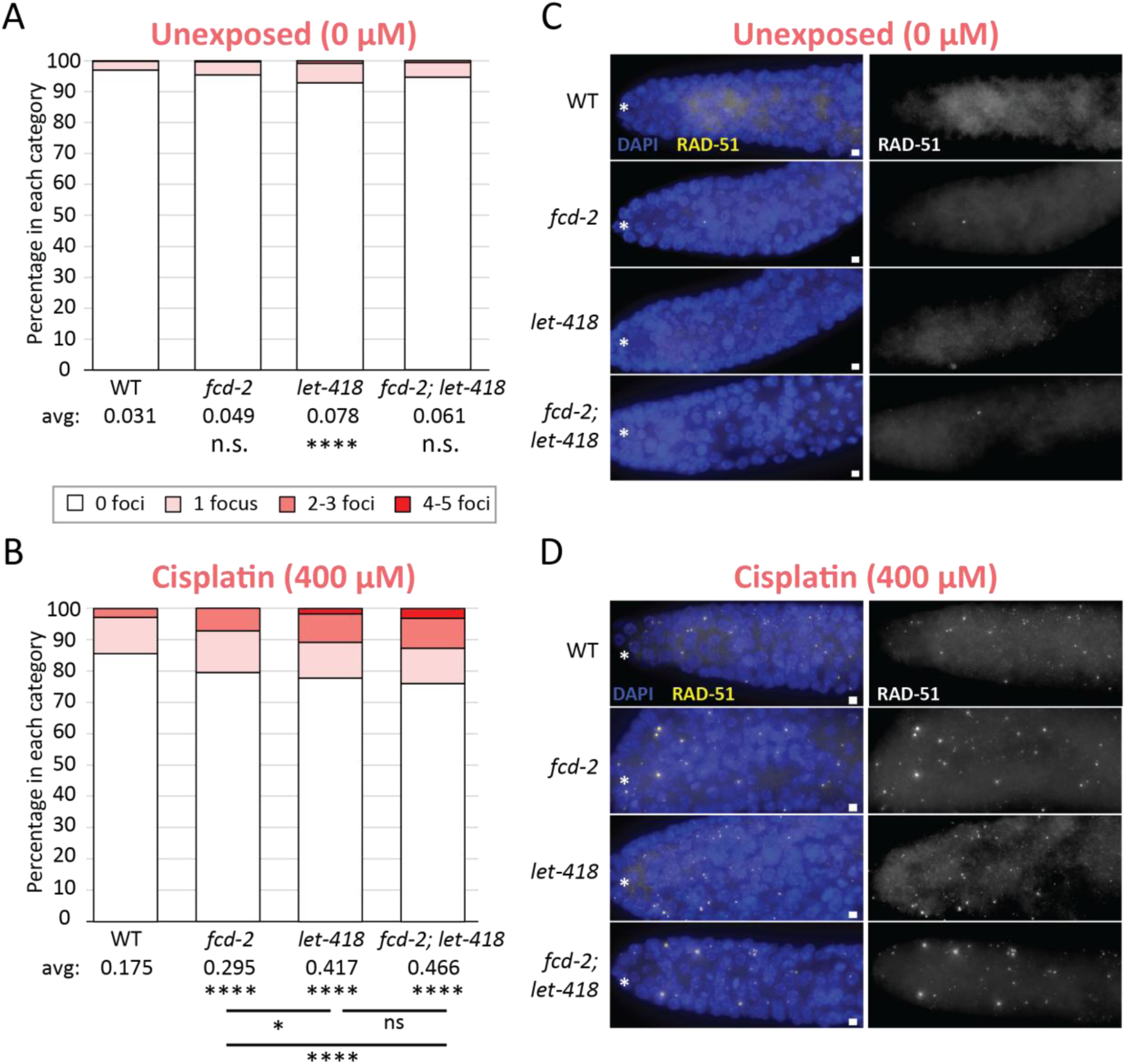
*let-418* mutants cannot repair mitotic DSBs when challenged with cisplatin. **(A,B)** Stacked histograms showing the percentage of nuclei with the indicated number of RAD-51 foci for animals exposed to 0 μM (A) and 400 μM of cisplatin (B). Average number of foci per nucleus is indicated under each genotype. Comparisons between wild-type and mutant populations at the same dose are shown directly under averages for each genotype; other comparisons are indicated by horizontal lines. For all genotypes, exposed populations differed significantly from unexposed controls (not indicated on histogram, *P* < 0.0001 for all comparisons). Each condition includes at least 2000 nuclei analyzed from 15 germlines across three biological replicates. (**C, D**) Representative images of mitotic region nuclei in unexposed (C) and exposed (D) animals, with the distal tip oriented to the left (as indicated by white asterisk). Germlines are stained for RAD-51 (in yellow) and DAPI (in blue). Scale bar represents 2 μm. All statistical analyses were performed using a Kruskal-Wallis test with Dunn’s post test (n.s. = not significant, * *P* < 0.05, **** *P* < 0.0001). Summary statistics and data are included in Table S1.

**Figure 2:**
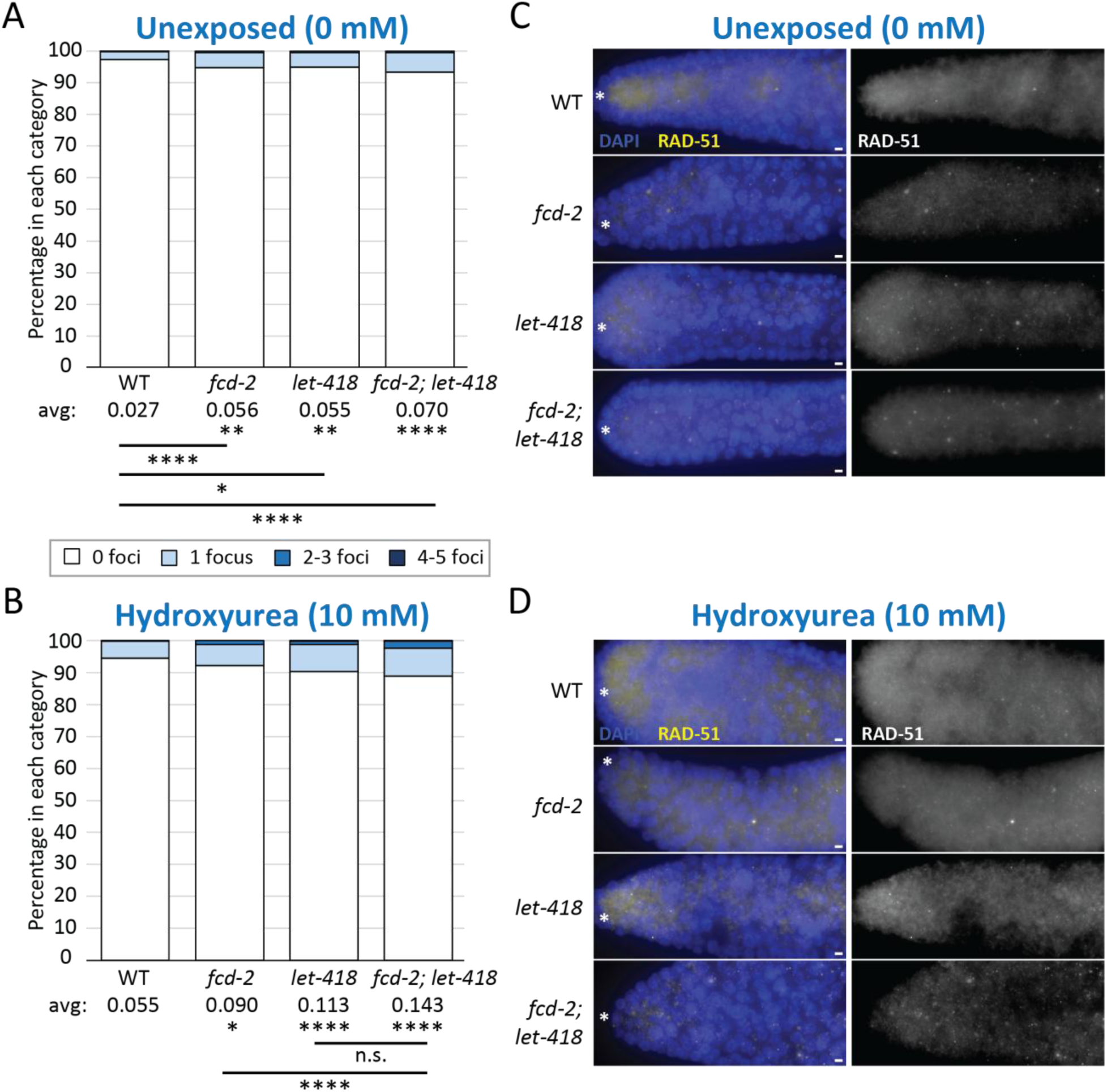
*let-418* mutants cannot repair mitotic DSBs when challenged with hydroxyurea. **(A, B)** Stacked histograms showing the percentage of nuclei with the indicated number of RAD-51 foci for animals exposed to 0 mM (A) and 10 mM of hydroxyrea (B). Average number of foci per nucleus is indicated under each genotype. Comparisons between wild-type and mutant populations at the same dose are shown directly under averages for each genotype; other comparisons are indicated by horizontal lines. For all genotypes, exposed populations differed significantly from unexposed populations (not shown on graph, *P* < 0.0001 for all comparisons). Each condition includes at least 2000 nuclei analyzed from 15 germlines across three biological replicates. (**C, D**) Representative images of mitotic region nuclei in unexposed (C) and exposed (D) animals, with the distal tip oriented to the left (as indicated by white asterisk). Germlines are stained for RAD-51 (in yellow) and DAPI (in blue). Scale bar represents 2 μm. All statistical analyses were performed using a Kruskal-Wallis test with Dunn’s post test (n.s. = not significant, * *P* < 0.05, ** *P* < 0.01, **** *P* < 0.0001). Summary statistics and data are included in Table S1.

As expected, exposure to 400 μM cisplatin increased the number of RAD-51 foci across all genotypes compared to unexposed controls (Fig. 1B, D and Table S1). This dose was chosen based on a dose curve showing that 400 μM drastically reduced embryonic survival in both wild-type and mutant populations (Fig. S1 and Table S6). Wild-type populations exposed to cisplatin accumulated over five times more foci than their unexposed controls *P* < 0.0001, Kruskal-Wallis test) (Fig. 1A, B). Similarly, both exposed *fcd-2* and *let-418* mutants accumulated five times more foci than unexposed controls (*P* < 0.0001 for both comparisons, Kruskal-Wallis test) (Fig. 1A, B). In *let-418* mutants, the increase was driven by nuclei with multiple foci, sometimes as many as five per nucleus (10.8% of nuclei compared to 0.7%). Interestingly, exposed *fcd-2; let-418* double mutants strongly resembled exposed *let-418* single mutants, with over seven times more foci than unexposed controls (*P* < 0.0001, Kruskal-Wallis test) (Fig. 1B). Again, this increase was primarily driven by nuclei with multiple foci, especially those with four or five foci (9.6% of nuclei compared to 0.7%). Notably, nuclei with four or five foci were never observed in exposed wild-type or *fcd-2* single mutants. All mutants examined were also more sensitive to cisplatin than wild-type controls. Exposed *fcd-2* single mutants experienced 1.7 times more foci than exposed wild-type populations (*P* < 0.0001), whereas exposed *let-418* single mutants were even more affected, with 2.4 times more foci than exposed wild-type populations (*P* < 0.0001 compared to WT, *P* < 0.05 compared to *fcd-2*). The increase in exposed *fcd-2; let-418* double mutants resembled that seen in *let-418* single mutants (*P* > 0.9), with the double mutants having 2.7 times more foci than exposed wild-type populations (*P* < 0.0001).

We next challenged mutants using hydroxyurea, which stalls replication forks and generates DSBs if left unrepaired (Bailly and Gartner, 2011). A dose of 10 mM was chosen after examining embryonic survival in wild-type and *let-418* mutants across a dose curve (Fig. S3 and Table S6). Hydroxyurea affected all genotypes in a manner similar to cisplatin (Fig. 2 and Table S1). Once again, few DSBs were observed in each unexposed population, with less than 7% of mitotic nuclei containing a RAD-51 focus (Fig. 2A). As expected, exposure to hydroxyurea raised the average number of foci in all genotypes (*P* < 0.02 for all comparisons, Kruskal-Wallis test) (Fig. 2B). Exposed wild-type populations and *fcd-2* mutants had nearly two times more foci than unexposed controls (*P* < 0.02 for both comparisons, Kruskal-Wallis test), both primarily driven by an increase in the number of nuclei with one focus. Resembling what we observed after cisplatin exposure, both single mutants were more sensitive to hydroxyurea than wild-type populations: exposed *fcd-2* single mutants had 1.6 times more foci than exposed wild-type controls (*P* < 0.05) and *let-418* single mutants had twice as many foci as exposed wild-type populations (*P* < 0.0001). As with cisplatin exposure, the response in *fcd-2; let-418* double mutants resembled the response in *let-418* single mutants (*P* > 0.9), with 2.6 times more foci than exposed wild-type populations (*P* < 0.0001). We note that exposed *let-418* and *fcd-2; let-418* mutants each contained nuclei with as many as five RAD-51 foci, which were almost never observed in exposed wild-type or *fcd-2* mutant populations (Fig. 2B).

To examine whether these unrepaired exogenous DSBs persisted from mitosis into meiosis, we assessed RAD-51 foci throughout pachytene (Fig. 3 and Table S2). Hermaphrodites were exposed to cisplatin or hydroxyurea for twenty hours, allowed to recover for 30-40 hours (to allow mitotic nuclei to progress into mid- or late-pachytene), and dissected for immunofluorescence (Jaramillo-Lambert et al., 2007; Kessler and Yanowitz, 2014). At mid-pachytene, wild-type animals had an average of 3.3 foci per nucleus, which is consistent with prior observations (Fig. 3A, B) (Achache et al., 2021; Cahoon et al., 2023; Colaiácovo et al., 2003; Germoglio et al., 2020; Rosu et al., 2013; Stamper et al., 2013). All unexposed mutants had significantly more RAD-51 foci than unexposed wild-type animals in mid-pachytene (*P* < 0.001 for all mutants, Kruskal-Wallis test; see Table S2 for additional comparisons between early-pachytene and late-pachytene), and the number of foci in *fcd-2; let-418* double mutants resembled that in both single mutants (*P* > 0.9) (Fig. 3A, B).

**Figure 3:**
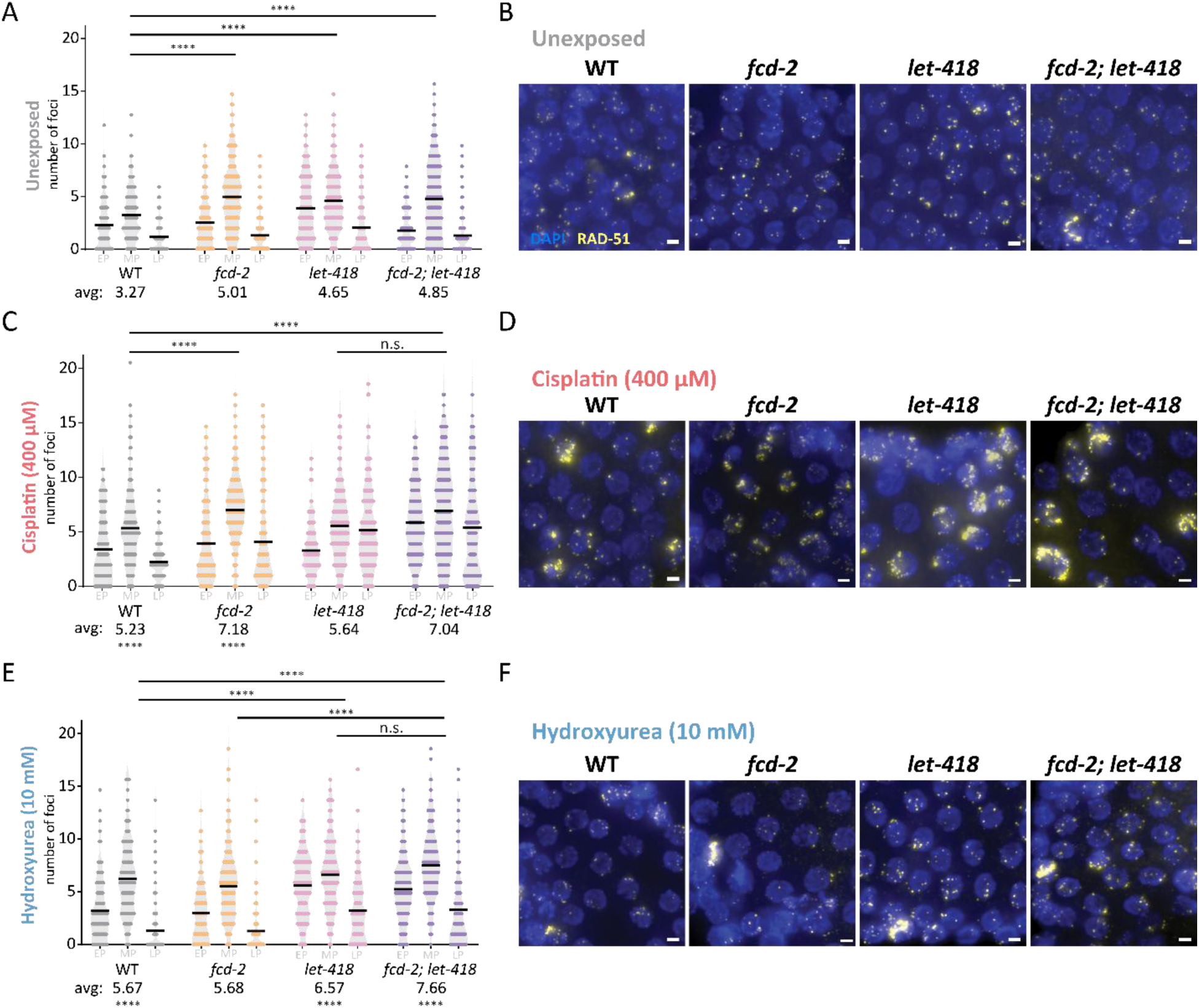
Mutagen-induced DNA damage persists throughout pachytene in *let-418* mutants. **(A, C, E)** Quantification of number of nuclei with the indicated number of RAD-51 foci in unexposed (A), cisplatin-exposed (C), and hydroxyurea-exposed (E) hermaphrodites. Dark line indicates average number of foci for each stage of pachytene: early pachytene (EP), mid-pachytene (MP), and late pachytene (LP). Each stage represents at least 100 nuclei scored from three gonads. Averages for mid-pachytene are shown under each genotype. All comparisons shown are only for mid-pachytene regions (comparisons for early- and late-pachytene regions are included in Table S2): comparison between exposed and unexposed populations are shown under averages; other comparisons are indicated by horizontal lines. **(B, D, F)** Representative immunofluorescence images of mid-pachytene nuclei stained for RAD-51 (yellow) and DAPI (blue) in unexposed (B), cisplatin-exposed (D), and hydroxyurea-exposed (F) hermaphrodites. Scale bar represents 2 μm. Summary statistics and data are included in Table S2. All statistical analyses were performed using a Kruskal-Wallis test with Dunn’s post test (n.s. = not significant, **** *P* < 0.0001).

Exposing mitotic nuclei to cisplatin significantly increased the number of foci during mid- or late-pachytene across all genotypes compared to unexposed controls (*P* < 0.0001 for all comparisons) (Fig. 3C, D). Notably, all mutants had persistent damage even in late-pachytene (Fig. 3C and Table S2) (*P* < 0.05 for all genotypes compared to wild-type) – both *let-418* single mutants and *fcd-2; let-418* double mutants were more affected than *fcd-2* single mutants, with each experiencing an average of over five foci per nucleus in late-pachytene (*P* < 0.05 for each compared to *fcd-2*, *P* > 0.9 for double mutant compared to *let-418*). Similarly, exposure to hydroxyurea significantly increased foci during mid-pachytene in both *let-418* single mutants and *fcd-2; let-418* double mutants compared to unexposed controls (*P* < 0.0001 for both comparisons) (Fig. 3E, F), and this increase persisted into late-pachytene when compared to exposed wild-type animals or *fcd-2* single mutants (*P* < 0.0001 for all comparisons) (Fig. 3E and Table S2).

Taken together, these data indicate that cisplatin or hydroxyurea generate mitotic DNA lesions that require both LET-418 and FCD-2 for proper repair. In their absence, exogenous damage incurred during mitosis persists until the end of pachytene. Additionally, when responding to mitotic DNA damage in the germline, *fcd-2; let-418* double mutants phenocopy *let-418* single mutant for both mutagens.

### Unrepaired mitotic DSBs affect oocyte quality in *let-418* mutants

Although *let-418* single mutants accumulate DSBs in the mitotic region and during pachytene, they do not display aneuploidy or chromosome fragmentation, indicating that any remaining DSBs are fully resolved by diakinesis (Turcotte et al., 2018). However, mutants lacking both CHD paralogs, CHD-3 and LET-418, suffer from meiotic catastrophe, genome fragmentation, and chromosome fusions (Turcotte et al., 2018), suggesting that NuRD needs at least one CHD component to repair endogenous DSBs. To see whether LET-418 is required for the resolution of exogenous DNA lesions, we exposed hermaphrodites to cisplatin or hydroxyurea for twenty hours and allowed them to recover for sixty hours (Jaramillo-Lambert et al., 2007; Kessler and Yanowitz, 2014). This timing allows us to induce DNA damage in mitotic cells and assess its impact during diakinesis, where we expect to see six DAPI-staining bodies, one for each homologous pair of chromosomes.

As expected, unexposed wild-type animals rarely produced aneuploid gametes: when scoring aged mothers in their third day of adulthood, we found that only 9% of oocytes contained five DAPI-stained bivalents (Fig. 4A, C, E, Fig. S2, and Table S3). This result is consistent with previous studies showing that oocyte quality decreases with maternal age (Achache et al., 2021; Andux and Ellis, 2008; Luo et al., 2010; Scharf et al., 2021). Each unexposed mutant produced more aneuploid oocytes than wild-type controls, with at least 16% of nuclei containing fewer than six DAPI bodies (*P* < 0.01 for all comparisons, Fisher’s exact test), including oocytes with four DAPI bodies that were never observed in wild-type animals (Fig. 4A, C, E and Fig. S2). We also noted that *let-418* single mutants generated a rare oocyte with seven DAPI bodies (Fig. 4E). The presence of univalents in this nucleus reflected the lack of a chiasmata and failure of crossover formation, which almost never occurs in wild-type animals (Dernburg et al., 1998; Yu et al., 2016). When exposed to 400 μM cisplatin, both *fcd-2* single mutants and *fcd-2; let-418* double mutants suffered a significant deterioration in oocyte quality (*P* < 0.04 for both comparisons, Fisher’s exact test) (Fig. 4B). Each mutant produced oocytes with more than six DAPI bodies, and *fcd-2; let-418* double mutants also produced an oocyte with only three DAPI bodies (Fig. 4F). Although exposure to cisplatin did not induce a statistically significant change in wild-type and *let-418* single mutants (*P* > 0.5 for both comparisons, Fisher’s exact test), both genotypes produced oocyte classes that were never observed in unexposed controls (Fig. 4E, F and Table S3).

**Figure 4:**
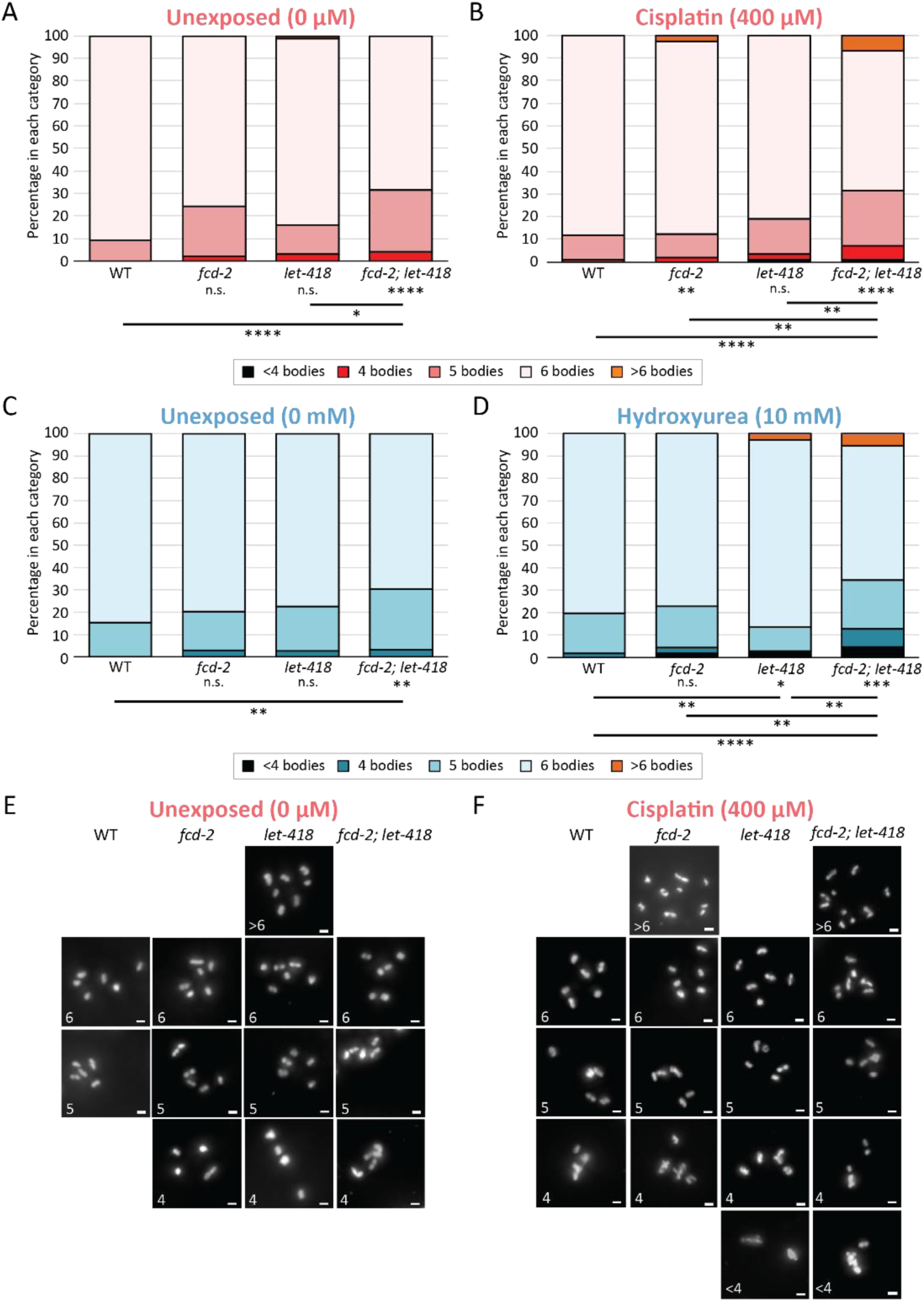
*let-418* mutants have poor oocyte quality due to mutagen induced mitotic DSBs. **(A, B, C, D)** Stacked histograms showing the percentage of nuclei with the indicated number of DAPI bodies for animals exposed to 0 μM of cisplatin (A), 400 μM of cisplatin (B), 0 mM of hydroxyurea (C) and 10 mM of hydroxyurea (D). Comparisons between wild-type and mutant populations at the same dose are shown directly under averages for each genotype; other comparisons are indicated by horizontal lines. Each condition includes at least 100 nuclei analyzed from 27 germlines across three biological replicates. **(E, F)** Representative images of -1 oocytes in animals exposed to 0 uM (E) and 400 uM of cisplatin (F). See Fig. S1 for representative images of -1 oocytes exposed to hydroxyurea. Germlines are stained with DAPI (in white). Scale bar represents 2 μm. All statistical analyses were performed as a Fisher’s exact test on number of DAPI bodies (n.s. = not significant, * *P* < 0.05, ** *P* < 0.01, **** *P* < 0.0001). Summary statistics and data are included in Table S3.

When compared to either single mutant, *fcd-2; let-418* double mutants were more affected by cisplatin exposure (*P* < 0.003 for both comparisons, Fisher’s exact test). This difference was driven by oocytes with both more and fewer DAPI bodies: *fcd-2; let-418* double mutants produced more nuclei with four or fewer DAPI bodies, but also nuclei with seven to ten DAPI bodies, indicating a severe disruption in crossover formation (Fig. 4F). Given our findings for mitotic DSBs, where the double mutant phenocopied the *let-418* single mutant (Fig. 1 and Fig. 2), we were surprised to see that diakinesis DAPI body number was significantly more affected in *fcd-2; let-418* double mutants than in *let-418* single mutants. However, both the double mutant and *let-418* single mutant produced the rare category of nuclei with three DAPI bodies (Fig. 4F). The severe reduction in DAPI bodies indicated that at least half of the *C. elegans* genome has experienced chromosomal fusions, a phenotype that was also previously observed in mutants missing both CHD paralogs (Turcotte et al., 2018).

Similar to cisplatin, exposure to 10 mM hydroxyurea also induced aneuploidy during diakinesis (Fig. 4C, D and Table S3). Although exposure to hydroxyurea did not cause a statistically significant change in wild-type population or *fcd-2* single mutants (*P* > 0.3 for both comparisons, Fisher’s exact test), both genotypes produced oocyte classes that were never observed in unexposed controls (Fig. S2 and Table S3). However, unlike what we observed after cisplatin exposure, we did not find that *fcd-2* single mutants produced nuclei with more than six DAPI bodies after hydroxyurea exposure (Fig. 4D), indicating that *fcd-2* mutants were more sensitive to cisplatin. Exposed *let-418* single mutants were significantly affected compared to wild-type controls, producing oocytes with both more than seven and fewer than four DAPI bodies, categories that were never observed in unexposed controls (*P* < 0.006, Fisher’s exact test).

Challenging *fcd-2; let-418* double mutants with hydroxyurea caused a large deterioration in oocyte quality (*P* < 0.009, Fisher’s exact test). Exposed *fcd-2; let-418* double mutants were significantly more affected than either of the exposed single mutant controls (*P* < 0.004 for both comparisons, Fisher’s exact test). This decline in oocyte quality was partly driven by nuclei with more than six DAPI bodies (sometimes as many as ten), a category that was only ever observed in exposed *fcd-2; let-418* double mutants and *let-418* single mutants (Fig. S2B and Table S3). Once again, the existence of nuclei with univalents indicates that hydroxyurea also causes a severe failure of crossover formation or chromosome fragmentation (Fig. S2B). Taken together, this assessment of diakinesis nuclei indicates that *fcd-2; let-418* double mutants are more sensitive to interstrand crosslinks and stalled replication forks than either single mutant.

### Mutagen exposure reduces embryonic survival of *let-418* mutants

After having shown that LET-418 is necessary to repair exogenous DSBs during mitosis and meiosis (Fig. 1 through 3) and prevent aneuploidy during diakinesis (Fig. 4), we next examined oocyte viability by assessing embryonic survival (Fig. 5 and Table S4). To capture the impact of mutagen exposure during mitosis, we assessed embryos 60 hours after exposure, which were produced by mothers in their third day of adulthood (Jaramillo-Lambert et al., 2007; Kessler and Yanowitz, 2014). As expected for aged mothers, unexposed wild-type populations had an average embryonic survival of 96% (Fig. 5A). As a positive control for cisplatin sensitivity, we examined *xpf-1 (tm2842)* mutants, which lack an endonuclease involved in the DNA damage response (Ward et al., 2007). Unexposed *xpf-1* mutants have a lower average survival than unexposed wild-type populations (*P* < 0.007, ANOVA) (Fig. 5A), which is reduced even further by cisplatin in a dose-dependent manner (Fig. S1B and Table S6) (Meier et al., 2014; Ward et al., 2007). Survival in unexposed *fcd-2*, *let-418*, or *fcd-2; let-418* mutants did not significantly differ from wild-type populations (*P* > 0.1 for all comparisons), although each mutant displayed more variability and lower averages across replicates compared to wild-type: across both cisplatin and hydroxyurea experiments, *fcd-2* single mutants averaged 86% survival, *let-418* single mutants averaged 77% survival, and *fcd-2; let-418* double mutants averaged 75% survival (Table S4). When challenged with 400 μM cisplatin, embryonic survival in wild-type populations was significantly reduced to 73% (*P* < 0.04). As expected for a positive control, cisplatin exposure reduced *xpf-1* mutant survival more severely than in wild-type populations, down to 39% survival (*P* < 0.01) (Fig. 5A).

**Figure 5:**
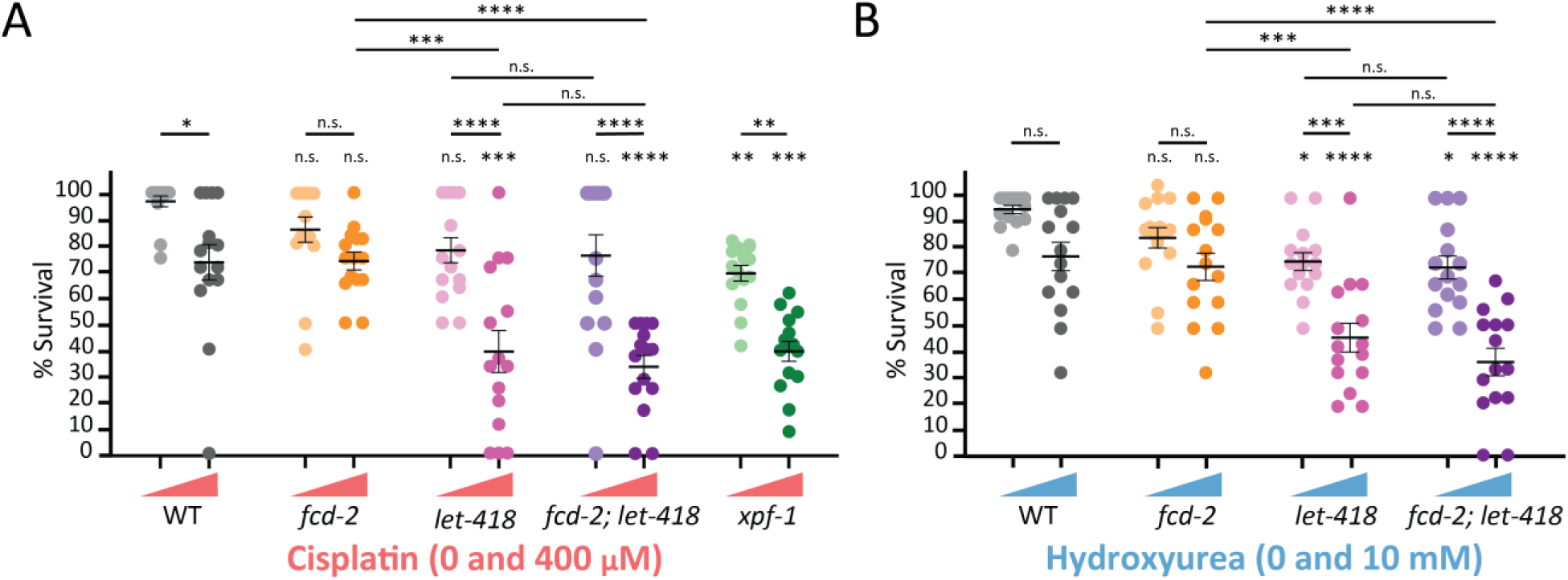
DNA damaging agents reduce embryonic viability in *let-418* mutants. (**A, B**) Percent survival of indicated genotypes with and without exposure to cisplatin (A) or hydroxyurea (B); dose is indicated with triangle above genotype names, with cisplatin in salmon (0 μM and 400 μM) and hydroxyurea in blue (0 mM and 10 mM). For all conditions in (A) N=125 and (B) N=150. Dark line represents the mean and whiskers represent S.E.M. Comparisons between wild-type and mutant populations at the same dose are above each condition; other comparisons are indicated by horizontal lines. All statistical analyses were performed using a one-way ANOVA with a Šídák correction (n.s. = not significant, \**P* < 0.05, ** *P* < 0.01, *** *P* < 0.001, **** *P* < 0.0001). Summary statistics and data are found in Table S4.

Embryonic survival in exposed *fcd-2* single mutants was not significantly affected compared to unexposed controls (*P* > 0.8, ANOVA) (Fig. 5A). Conversely, exposed *let-418* single mutants experienced a significant decline in survival, down to 39% (*P* < 0.0001), a decrease 2.4 times more than wild-type and similar to *xpf-1* mutants (Fig. 5A). We noted that cisplatin had a variable impact on *let-418* mutant survival, across both technical and biological replicates. This observation was consistent with previous variability observed in *let-418* mutants (Turcotte et al., 2018): some replicates were completely unaffected, resembling unexposed controls, whereas others suffered complete embryonic lethality that was never observed in wild-type or *xpf-1* positive controls. Again, the effects of cisplatin on embryonic survival in *fcd-2; let-418* double mutants strongly resembled *let-418* single mutants (Fig. 5A). Exposed *fcd-2; let-418* double mutants experienced a significant drop in survival down to 33% (*P* < 0.0001). As seen in *let-418* single mutants, this reduction was 2.7 times more than wild-type and similar to *xpf-1* positive controls. However, in one departure from *let-418* single mutants, we found that cisplatin caused less variability in *fcd-2; let-418* double mutant survival, which had a distribution that overlapped only the lower half of that observed in *let-418* single mutants (Fig. 5A).

We next challenged hermaphrodites with 10 mM hydroxyurea to assess the impact on embryonic survival (Fig. 5B). As expected, this low dose of exposure did not significantly reduce survival in wild-type populations or *fcd-2* single mutants (*P* > 0.09 for both comparisons, ANOVA) (Kim et al., 2018). However, *let-418* single mutants were sensitive to hydroxyurea – embryonic survival dropped from 76% in unexposed controls to 46% after exposure (*P* < 0.0003), a reduction 2.1 times more than experienced by wild-type populations (Fig. 5B). Similarly, survival in exposed *fcd-2; let-418* double mutants dropped from 73% down to 36% (*P* < 0.0001), a reduction 2.8 times more than wild-type (*P* < 0.0001) (Fig. 5B).

Finally, we challenged hermaphrodites with an additional mutagen, nitrogen mustard, which creates both interstrand crosslinks and DNA-protein crosslinks (Fig. S4 and Table S6) (Povirk and Shuker, 1994). A low dose of 100 μM did not affect wild-type embryonic survival (*P* > 0.9, ANOVA), whereas a higher dose of 150 μM significantly decreased wild-type survival to 52% (*P* < 0.0001) (Fig. S4). The low 100 μM dose of nitrogen mustard decreased survival in both *let-418* and *fcd-2* single mutants (*P* < 0.0001 for both comparisons) and the high 150 μM dose further reduced survival to 25% (*P* < 0.0001 for both comparisons, ANOVA). Nitrogen mustard affected *fcd-2; let-418* double mutants similarly to each single mutant (*P* > 0.6 for all comparisons), with survival severely reduced to 10% by the high 150 μM dose (*P* < 0.0001 compared to unexposed control) (Fig. S4). Taken together, the impact of mutagen exposure on embryonic survival indicate that LET-418 is required to resolve exogenous DNA lesions to maintain oocyte viability.

### LET-418’s ATPase domain is necessary for its role in germline DSB repair

The protein sequences of *C. elegans* LET-418 and human CHD4 share extensive homology, including three domains: an ATPase domain, a PHD finger domain, and a double chromodomain (Fig. 6A) (Farnung et al., 2020; Käser-Pébernard et al., 2016; Passannante et al., 2010). Computational models of CHD4 structure suggest that each domain has distinct functions: the ATPase domain slides nucleosomes along DNA (Farnung et al., 2020), the PHD finger domains binds specific residues on histone H3 (Farnung et al., 2020; Mansfield et al., 2011) and the chromodomains primarily bind DNA (Bouazoune et al., 2002). Mutations discovered in human patients, along with *in vitro* experiments of CHD4 activity, have implicated each of these domains in CHD4’s nucleosome remodeling activity (Farnung et al., 2020; Watson et al., 2012; Weiss et al., 2016), but the impact of each domain has yet to be assessed *in vivo*. Because the kinetics of repair occur rapidly in response to DNA lesions (Kochan et al., 2017; Nair et al., 2017), we hypothesized that LET-418’s function in the germline requires its ATPase domain to slide nucleosomes and expose the site of DNA damage.

**Figure 6:**
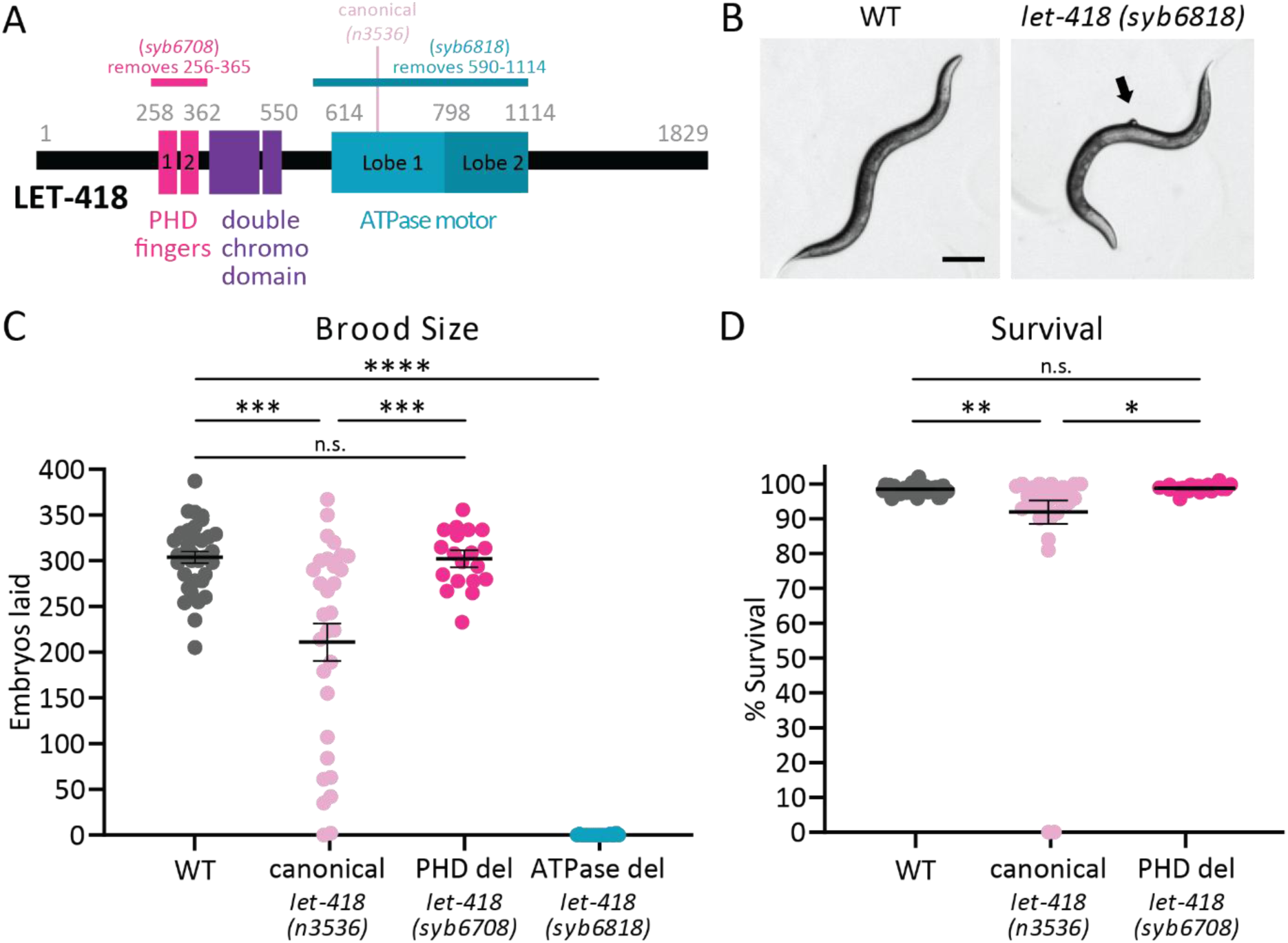
The ATPase motor domain is necessary for LET-418 protein function in DSB repair. **(A)** Protein alignment of human CHD4 and *C. elegans* LET-418 homologs showing conserved domains. Primary structural alignments were performed according to NCBI-predicted domains using UniProt. Locations of each allele are indicated above the alignment: *syb6708* PHD finger deletion (magenta), canonical *n3536* missense mutation (pink), and *syb6818* ATPase deletion (teal). (**B**) Images of young adult hermaphrodites. The ATPase deletion *syb6818*, has a protruding vulva phenotype (arrow to pVulv) and a lack of germline as indicated by the white space in the body. Scale bar, 100 µm. (**C**) Total brood was assessed in wild-type (gray), *n3536* canonical allele mutants (light pink), *syb6708* PHD finger deletion mutants (magenta), and *syb6818* ATPase deletion mutants (teal). (**D**) Percent embryonic survival, excluding *syb6818* due to complete sterility. For both graphs, dark line represents mean and whiskers represent S.E.M. All statistical analyses were performed using ANOVA with a Šídák correction (n.s. = not significant, \**P* < 0.05, \*\**P* < 0.01, *** *P* < 0.001, **** *P* < 0.0001). Summary statistics and data are included in Table S5.

Figures 1 through 5 of this paper relied on the canonical allele *let-418 (n3536)*, a point mutation in the ATPase domain that is hypomorphic, but viable, at the normal husbandry temperature of 20°C (Andersen et al., 2006; Turcotte et al., 2018). We engineered deletion alleles in the *let-418* gene using CRISPR-Cas9 to remove either the PHD finger domain or the ATPase enzymatic domain (see Methods for details). We then assessed the number of embryos produced throughout the entire laying period in each *let-418* mutant (a full brood) under normal husbandry conditions, using the canonical *n3536* allele as a positive control (Fig. 6A and Table S5). Consistent with previous studies, wild-type mothers produced broods that averaged 311 embryos (Fig 6C). Strikingly, mutants that lack the ATPase domain*, syb6818*, were completely sterile and produced no embryos (Fig. 6C). In these ATPase deletion mutants, we also noted a high occurrence of protruding vulvae and a clear body due to absence of germline, which accounts for the complete sterility (Fig. 6B). Conversely, mutants that lack the PHD finger domain, *syb6708*, produced an average brood of 304 embryos, which was not significantly lower than wild-type broods (*P* > 0.7, ANOVA). Consistent with previous reports, average broods of canonical *n3636* mutants were 188 embryos, which represents a significant reduction compared to wild-type broods (*P* < 0.001). We also noted the variability in canonical *n3636* mutant broods: some mothers produced at wild-type levels (∼300 embryos), whereas others were completely sterile (McMurchy et al., 2017; Turcotte et al., 2018).

Because LET-418 is also involved in development (Erdelyi et al., 2017; Passannante et al., 2010; von Zelewsky et al., 2000), we assessed embryonic survival throughout the full laying period. Average survival of canonical *n3636* mutants was 92%, which represented a slight but significant reduction compared to average wild-type survival of 98% (*P* < 0.01, ANOVA). In *syb6708* PHD deletion mutants, embryonic survival resembled wild-type survival at 99%, similar to what we observed with brood size (*P* > 0.9) (Fig. 6D). Finally, we were unable to assess survival in *syb6818* ATPase deletion mutants, because all mothers examined were sterile (N = 20). Altogether, our findings support a model in which LET-418’s ATPase domain, but not its PHD domain, is necessary for NuRD’s role during oogenesis in otherwise wild-type conditions. Future studies will examine whether the PHD domain is necessary for repairing exogenous DNA damage.

## DISCUSSION

The germline must preserve genome integrity for all future generations, a task that requires extensive coordination between DNA repair machinery and local chromatin environments (Chen and Tyler, 2022; Clouaire and Legube, 2019). At the site of DNA damage, chromatin landscapes help orchestrate repair using either error-prone or error-free DSB repair pathways (Chen and Tyler, 2022). Although the mechanisms of DNA repair have been well-described across species, the role of chromatin remodelers remains poorly understood. In this study, we show that LET-418, the ATPase remodeling subunit of NuRD, is required to maintain oocyte quality when genomes are challenged by exogenous DNA damage. After exposure to cisplatin or hydroxyurea, mutants with reduced LET-418 activity accumulate more DSBs in the germline (Figs. 1-3), generate more aneuploid oocytes (Fig. 4), and suffer from reduced embryonic viability (Fig. 5 and Fig. 6) – phenotypes that strongly resemble those seen in mutants defective for the FA repair pathway (Adamo et al., 2010; Gartner and Engebrecht, 2022; Lee et al., 2010). When testing the genetic relationship between NuRD and the FA pathway, we showed that *let-418* gene activity is epistatic to the FA gene *fcd-2*, which suggests that NuRD nucleosome remodeling is necessary to allow proper functioning of the FA repair pathway.

LET-418/CHD4 has been implicated in biological functions that include maintaining repressive chromatin states, repairing DNA damage, and defining cell fate during embryonic development (De Vaux et al., 2013; Guerry et al., 2007; Käser-Pébernard et al., 2016; Kunert et al., 2009; Saudenova and Wicky, 2018; Turcotte et al., 2018; Unhavaithaya et al., 2002; von Zelewsky et al., 2000). In mammals, CHD4’s role in resisting DNA damage has led to its upregulation in multiple types of cancer: for example, in glioblastoma, CHD4 overexpression is associated with poor prognosis, likely due to its role in promoting DNA damage repair to assure cancer cell survival (Chudnovsky et al., 2014; McKenzie et al., 2019). Other work in ovarian cancer cells, acute myeloid leukemia cells, and mammary cells has shown that depleting CHD4 impairs homologous recombination and sensitizes tumorigenic cells to DNA damaging agents (Pan et al., 2012; Polo et al., 2010; Sperlazza et al., 2015). Similarly, in *C. elegans*, mutations in LET-418 or its paralog CHD3 prevent DSB repair throughout meiotic pachytene, suggesting that LET-418/CHD4’s role is highly conserved across taxa (Turcotte et al., 2018).

Our work identifies a new role for LET-418/CHD4 in response to genotoxic threats – in the absence of LET-418’s ATPase activity, mutants cannot fully repair exogenous DNA damage incurred during mitosis, which ultimately reduces gamete viability and impairs organismal survival. In germlines, LET-418 is associated with other heterochromatin factors that work with small RNA pathways to silence repetitive elements and protect the genome (McMurchy et al., 2017). The DSB repair defects observed in *let-418* mutants may be caused by disruptions in heterochromatin formation or maintenance, leading to higher sensitivity to DNA damaging agents. A similar effect has been observed with the loss of another core component of NuRD, RBBP4 (LIN-53 in *C. elegans*), which sensitizes cells to DNA damaging agents by establishing a permissive chromatin environment and downregulating RAD51 expression (Kitange et al., 2016).

We propose that LET-418 and NuRD are required during at least two points in gametogenesis: to repair exogenous damage accumulated in mitosis and to resolve endogenous DSBs formed during meiosis (this study and Turcotte et al., 2018). In the absence of LET-418 or NuRD activity, mitotic damage persists as nuclei enter meiosis, contributing to the increase of DSBs observed in pachytene (this study and Turcotte et al., 2018). The accumulated damage persists until the end of pachytene, and is reflected in aberrant diakinesis oocytes. The increase in meiotic DSBs may be caused by an inability of the FA pathway to access the damage. During meiosis, the FA pathway continues to repair DNA damage caused by replication fork stalling or interstrand crosslinks – after the lesion is recognized, FA component FANCD2 is mono-ubiquitinated and recruited to the lesion, where it mediates a conversion to DSBs and repair by homologous recombination (Ceccaldi et al., 2016; Niraj et al., 2019). Therefore, the loss of LET-418 or NuRD activity may interfere with recognition of the lesion or with recruiting FCD-2 to the site of damage.

Cytological experiments show that FA components, including FCD-2/FANCD2, are recruited to DNA lesions to mediate repair in the mitotic region of the germline (Collis et al., 2006; Kim et al., 2018; Lee et al., 2010). Our results are consistent with FCD-2 playing a role in resolving mitotic DSBs – when exposed to mutagens, *fcd-2* single mutants generated aneuploid oocytes that were never observed in wild-type populations: some had chromosome fusions, whereas others experienced crossover failure (Fig. 4 and Fig. S2). These defects represent genomic instability that is a consistent hallmark of defects with the FA pathway, and is even used to diagnose human patients for Fanconi anemia (Adamo et al., 2010, 2010; Auerbach, 2009; De Winter and Joenje, 2009; Lee et al., 2007; Rageul and Kim, 2020). However, we were surprised to find that embryonic viability in *fcd-2* mutants was not affected by exposure to either cisplatin or hydroxyurea (Fig. 5), since others have previously reported a cisplatin sensitivity in this mutant background (Adamo et al., 2010; Collis et al., 2006; Germoglio et al., 2020). After examining the timing of exposure and recovery used in those studies, we realized that they primarily assessed how damage induced during late pachytene affected embryonic survival (Jaramillo-Lambert et al., 2007; Kessler and Yanowitz, 2014). In contrast, we used a time course that allowed us to examine the effects of damage induced earlier, during mitosis. Altogether, our results suggest that FCD-2 and the FA pathway are involved in repairing exogenous DSBs, but may not be the sole, or even the main, repair pathway at this point in gametogenesis. However, in the later stages of meiotic prophase, all DSBs must be resolved before chromosome segregation. Therefore, as nuclei approach diplotene, they rely on alternative error-prone DSB repair pathways like NHEJ to resolve remaining DSBs (Macaisne et al., 2018; Smolikov et al., 2007). These pathways usually result in non-crossovers and disruption of chiasmata formation, or can generate chromosome translocations or fusions (Gartner and Engebrecht, 2022); we observed both outcomes in diakinesis nuclei of *let-418* mutants.

We have also demonstrated that LET-418’s activity is epistatic to that of FCD-2 and the FA pathway – the *fcd-2; let-418* double mutant phenocopies the *let-418* single mutant when challenged by exogenous DNA damage in its repair of DSBs and embryonic survival (Fig. 1-3, and Fig. 5). When examining how mitotic DNA lesions affect diakinesis DAPI body formation, we observed that *fcd-2; let-418* double mutants were more severely affected by cisplatin and hydroxyurea than either single mutant (Fig. 4 and Fig. S2). Strikingly, double mutant germlines contained classes of diakinesis nuclei that were only seen in one or the other single mutant, suggesting an additive effect of losing both LET-418 and FCD-2 activity. This result indicates that the FA pathway and NuRD may function in separate DSB repair pathways. The use of distinct pathways may be influenced by the region of the germline in which damage was incurred – in addition to the exogenous damage in mitosis caused by mutagen exposure, *let-418* mutants also accumulate damage endogenous DSBs during pachytene, which are not further exacerbated by the loss of FCD-2 in unexposed double mutants (Fig. 4A, C) (Turcotte et al., 2018).

*C. elegans* LET-418 shares extensive homology with human CHD4, including an ATPase domain, a PHD finger domain, and a double chromodomain (Käser-Pébernard et al., 2016; Passannante et al., 2010). Computational models indicate that the ATPase domain repositions nucleosomes by sliding them along DNA (Farnung et al., 2020). When examining human patient mutations identified in cancer or the neurodevelopmental disorder Sifrim-Hitz-Weiss syndrome, we noticed that many missense mutations occurred in the ATPase domain, indicating its importance for LET-418 and NuRD function. (Farnung et al., 2020). To test the requirement for LET-418’s ATPase activity *in vivo*, we engineered an endogenous deletion of the ATPase domain (allele *syb6818*) and demonstrated that it is essential for LET-418’s function in the germline. ATPase deletion mutants were completely sterile, and all homozygote mutants had defective vulvas (Fig. 6). Conversely, deletion of the PHD domain did not affect fertility or embryonic survival, indicating that this domain is dispensable for LET-418’s role in the germline in the absence of exogenous DNA damage. Our findings are consistent with deletion of the PHD domain in other contexts. In biochemical assays using recombinant human CHD4, deletion of the PHD domain causes a minor decrease in remodeling, but we note that this study used unmodified histones (Watson et al., 2012). In *Drosophila*, a hypomorphic point mutation to the PHD domain also slightly reduced remodeling ability, but did not otherwise affect binding of Mi2/CHD4 to chromatin (Kovač et al., 2018).

The ATPase deletion phenotypes resemble those of *let-418* genetic null alleles, which were previously used to characterize LET-418’s role in vulval development (Guerry et al., 2007; von Zelewsky et al., 2000). In somatic tissues, LET-418 belongs to two distinct complexes that have a conserved role in maintaining cell identity – in addition to NuRD, it is also found in MEC, which consists of LET-418, the histone deacetylase HDA-1, and the Krüppel-like protein MEP-1 (Hou et al., 2020; Käser-Pébernard et al., 2016; Pfefferli et al., 2014; Unhavaithaya et al., 2002). As part of these complexes, LET-418 has roles in specifying vulval fate, embryonic and larval development, and defining lifespan (De Vaux et al., 2013; Erdelyi et al., 2017; Guerry et al., 2007; Käser-Pébernard et al., 2014; Passannante et al., 2010; Saudenova and Wicky, 2018; von Zelewsky et al., 2000). Further characterization of these deletion strains will establish their impact after exposure to DNA damaging agents in in roles outside of the germline.

Based on our results, we propose the following model. When replication forks are blocked or stalled, NuRD’s remodeling activity creates a permissive chromatin environment that allows for the recruitment of FA pathway components, like FCD-2, to repair the damage via homologous recombination. In the absence of NuRD-mediated remodeling, the FA pathway is unable to repair DNA lesions, causing the accumulation of mitotic and meiotic DSBs and their eventual resolution by error-prone pathways later in meiosis. However, it is not clear from our findings whether NuRD functions entirely upstream of the full FA pathway, or whether it mediates an intermediate step during interstrand crosslink repair. For example, it is possible that DNA lesions are first recognized by the FA component FNCM-1 (which acts upstream of FCD-2), which then recruits NuRD to the site of damage. In summary, we’ve shown that LET-418’s nucleosome remodeling activity is necessary to resolve germline DSBs by error-free pathways like the FA pathway. This work highlights the important role of local chromatin landscapes in coordinating DNA damage repair pathways during gametogenesis.

## DATA AVAILIBILITY

Strains are available from the *Caenorhabditis* Genetics Center or upon request. The authors affirm that all data necessary for confirming the conclusions of the article are present within the article, figures, and tables.

## ACKNOWLEDGEMENTS

We thank Diana Libuda and Monica Colaiacovo for generously sharing reagents, Cori Cahoon for immunofluorescence advice, and WormBase for gene and sequence information (Davis et al., 2022). We also thank two anonymous reviewers for suggestions that strengthened the manuscript. Strains were provided by the *Caenorhabditis* Genetics Center, which is funded by the National Institutes of Health (NIH) through the Office of Research Infrastructure Programs (P40 OD010440).

## FUNDING

D. A. was supported by a UMass Lowell KCS Science Scholarship. T. B. and K. F. were supported by fellowships from the UMass Lowell River Hawk Scholars Academy. K. F. was supported by the Urban Massachusetts LSAMP program, which is funded by the National Science Foundation (EES 2308724). T.B. was supported by the Society for Developmental Biology through a Choose Development! fellowship, which is funded by the National Institutes of Health (R25HD105600). This work was supported by the National Institutes of Health, with grants R15GM117479 and R15HD104115 to P.M.C. and grant R15GM144861 to T.W.L.

## CONFLICT OF INTEREST

The authors declare no conflicts of interest.

---

**Figure S1:**
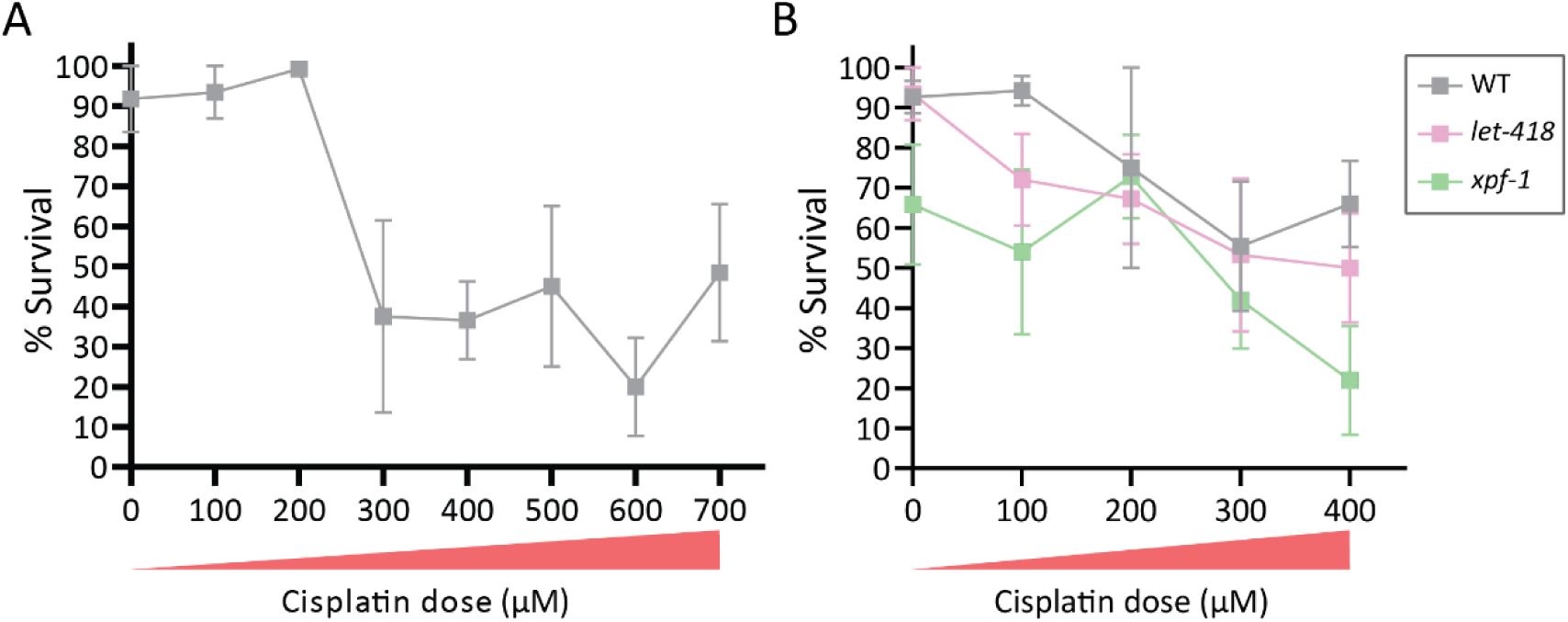
Cisplatin affects embryonic survival in a dose-dependent manner. (**A, B**) Mean percent survival of wild-type populations (gray) exposed to doses of cisplatin ranging from 0 μM to 700 μM (A) or 0 μM to 400 μM (B). Relative dose levels are indicated by triangles under the X-axis. (B) Examining the effects of cisplatin on *let-418* mutants (pink), with *xpf-1* mutants (green) included as a positive control for cisplatin sensitivity. Whiskers represent S.E.M. For each condition, at least 25 broods were evaluated. Summary statistics and data are included in Table S6.

**Figure S2:**
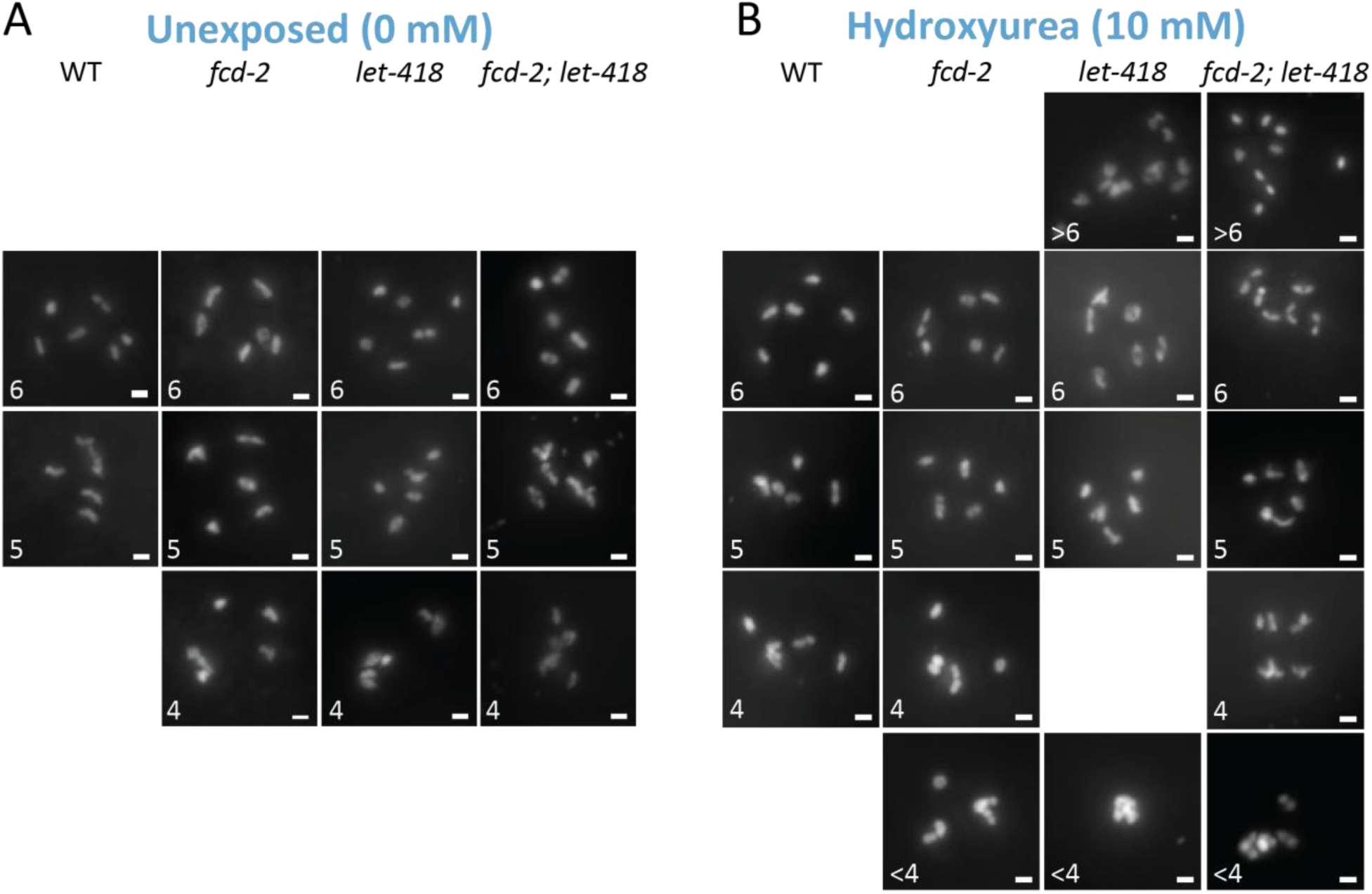
*let-418* mutants have poor oocyte quality due to hydroxyurea induced mitotic DSBs. (**A, B**) Representative images of -1 oocytes in unexposed (A) and exposed (B) animals. Germlines are stained with DAPI (in white). Scale bar represents 2 μm. Summary statistics and data are included in Table S3.

**Figure S3:**
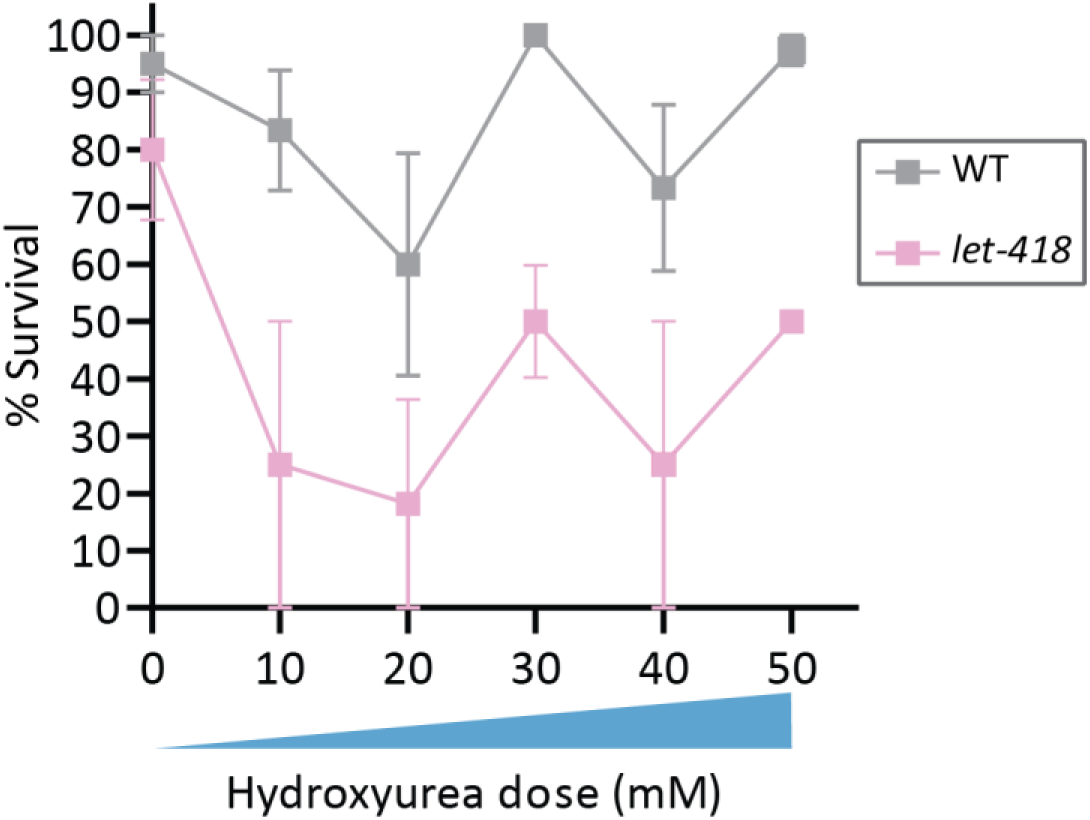
Hydroxyurea affects embryonic survival in a dose-dependent manner. Mean percent survival of wild-type (gray) and *let-418* mutants (pink) exposed to increasing doses of hydroxyurea (from 0 mM to 10 mM, with relative levels indicated by the blue triangle under the X-axis). Whiskers represent S.E.M. For each condition, at least 25 broods were evaluated. Summary statistics and data are included in Table S6.

**Figure S4:**
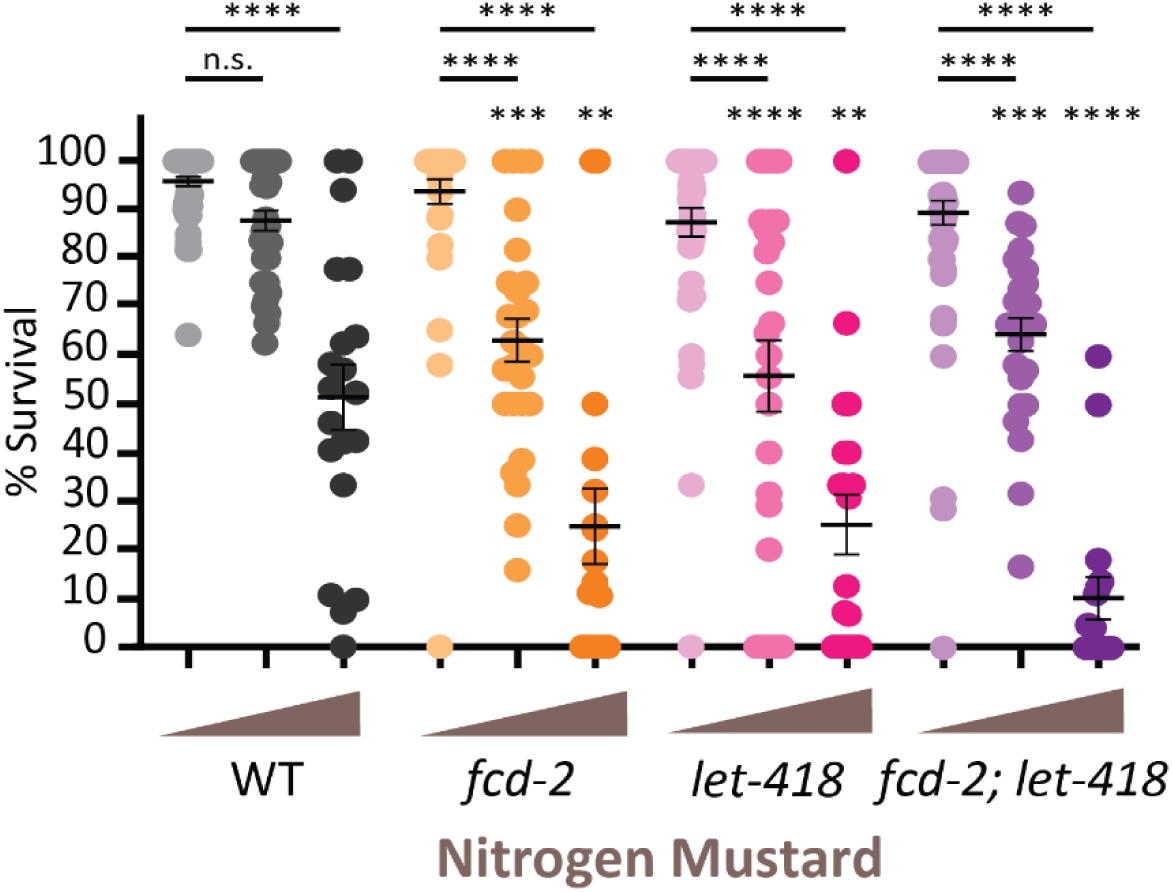
Nitrogen mustard reduces embryonic survival in *let-418* and *fcd-2* mutants. Percent survival of indicated genotypes with and without exposure nitrogen mustard; brown triangles indicate dose (0 mM, 100 mM, and 150 mM). Dark line represents mean percent survival and whiskers represent S.E.M. . Comparisons between wild-type and mutant populations at the same dose are above each mutant; other comparisons are indicated by horizontal lines. All statistical analyses were performed using ANOVA followed by Šídák’s multiple comparisons (\*\**P* < 0.01, *** *P* < 0.001, **** *P* < 0.0001). Summary statistics and data are included in Table S6.

## Notes

### Competing Interest Statement

The authors have declared no competing interest.

### Summary of Updates

New experiments examining DSBs in pachytene nuclei, updated figures, updated text.

